# Structural Heterogeneity of the Membrane-Interacting Region of the HIV-1 Envelope Glycoprotein

**DOI:** 10.1101/2025.09.16.676619

**Authors:** Ayan Majumder, Gregory A. Voth

## Abstract

The HIV-1 envelope glycoprotein (Env) trimer (gp120/gp41)_3_ forms the key functional envelope spike and is the target of neutralizing antibodies. The glycoprotein gp41 component mediates the fusion of viral and host cell membranes. In addition to its ectodomain, the membrane-interacting C-terminal domain of gp41 plays a crucial role in maintaining the fusogenicity and antigenic characteristics of Env. The membrane-interacting domain of gp41 consists of the highly conserved membrane proximal external region (MPER), which contains epitopes for broadly neutralizing antibodies, the transmembrane domain (TMD), which anchors Env in the membrane and mediates trimer formation, and the cytoplasmic tail (CT) domain, which plays an important role in Env trafficking to HIV-1 assembly sites. Previous experimental studies have extensively characterized the structure of the C-terminal domain of gp41; however, they reported different conformational states of the MPER and TMD. In this study, we used all-atom molecular dynamics simulations to investigate the structure and function of the membrane-interacting domain of gp41 in an HIV-1 mimetic membrane bilayer. The basic residues in the CT domain were found to interact favorably with PIP2, leading to lateral demixing of lipids and the accumulation of PIP2 in the cytofacial leaflet around the CT baseplate. Additionally, analysis based on an artificial intelligence (AI) machine learning based protocol revealed a diverse conformational ensemble of MPER-TMD, consistent with previous experimental observations. The MPER-TMD adopts both helix-turn-helix and extended helical conformations. We propose that the inherent flexibility of the MPER and the N-terminal region of TMD can play an important role in facilitating the late stages of membrane fusion.

**TOC Graphic:** 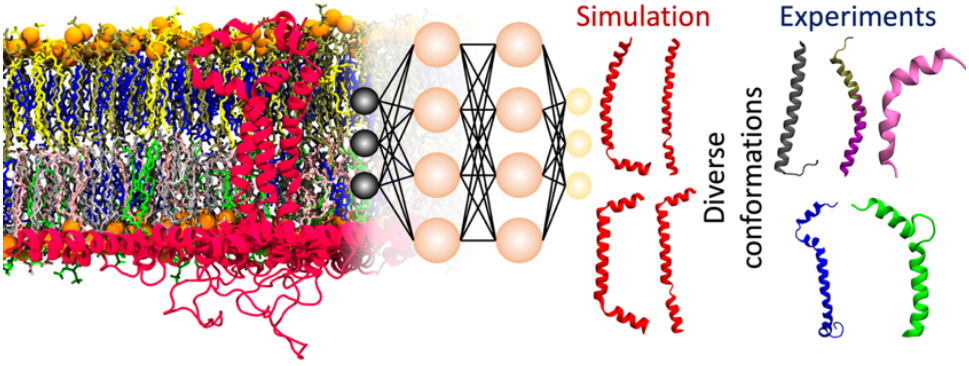

## Introduction

Human immunodeficiency virus type-1 (HIV-1) continues to infect people around the world, with no vaccine currently available.^1^ Since the beginning of the epidemic, HIV-1 has infected approximately 88 million people and caused nearly 42 million deaths.^2^ The envelope glycoprotein (Env) on the surface of the HIV-1 virion mediates entry into the host cell^3, 4^ and is the sole target of neutralizing antibodies.^5, 6^ Env is also an important target for the HIV-1 vaccine design efforts.^7^ Env is initially translated as a gp160 precursor protein containing about 850 residues. Gp160 trimerizes and is then cleaved by a furin protease into gp120 and gp41, which are primarily responsible for host cell receptor binding and membrane fusion, respectively.^8^ A trimer of the gp120-gp41 heterodimer forms a functional envelope spike on the surface of the HIV-1 virion.^9, 10^ This spike anchors to the viral membrane through the membrane proximal external region (MPER), transmembrane domain (TMD), and cytoplasmic tail (CT) domain of gp41 (**Figure 1**).^11^

**Figure 1:**
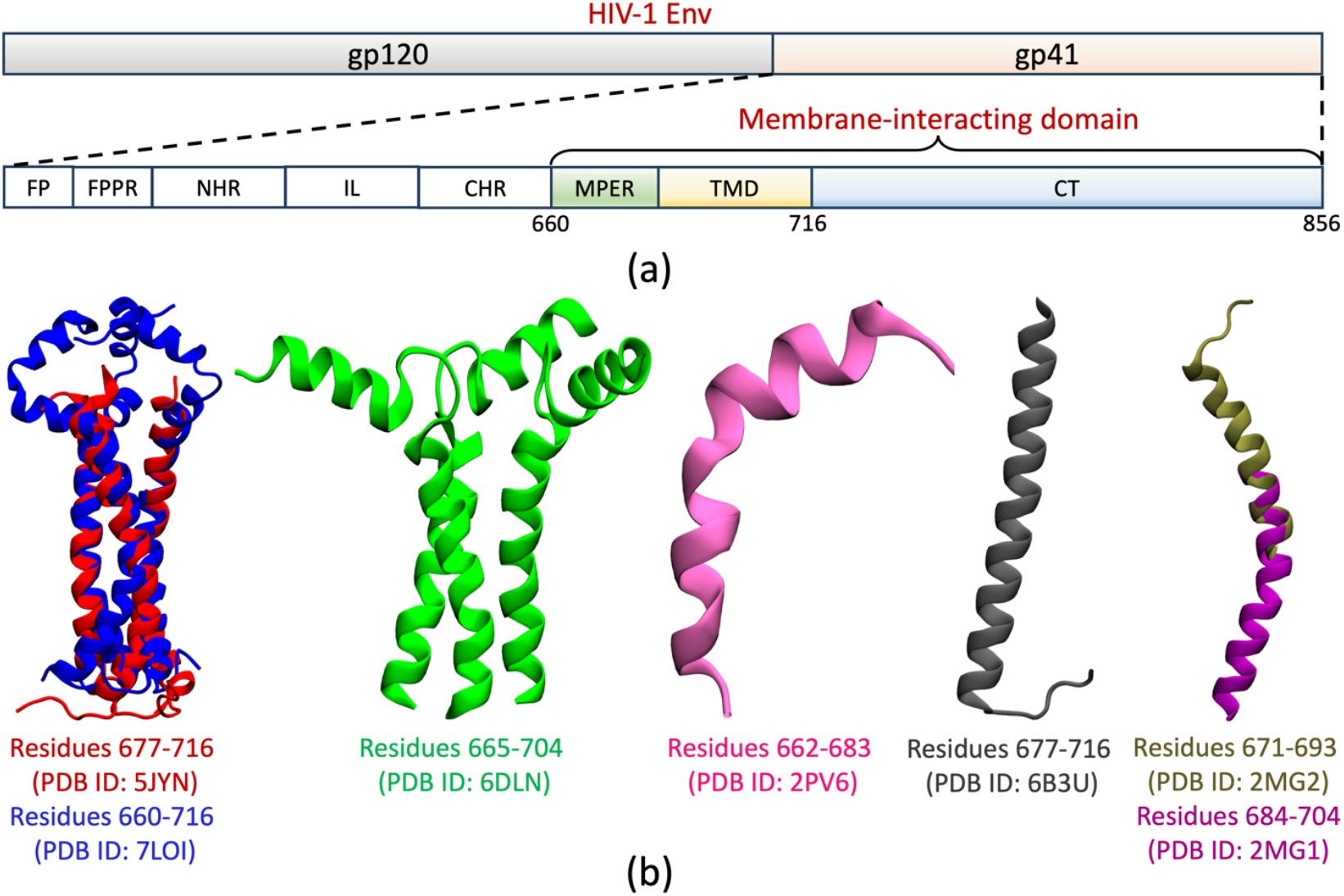
(a) Functional domains in gp41. The membrane-interacting domain of gp41 is simulated in this study. (b) Structures of the MPER-TMD region of gp41 reported by Chou and coworkers (red and blue)^11, 20^, Hong and coworkers (green)^24^, Reinherz and coworkers (pink)^26^, Bax and coworkers (gray)^25^, and Nieva and coworkers (tan and magenta)^27^. Experimental studies have revealed different conformations of the MPER-TMD.

**Figure 2:**
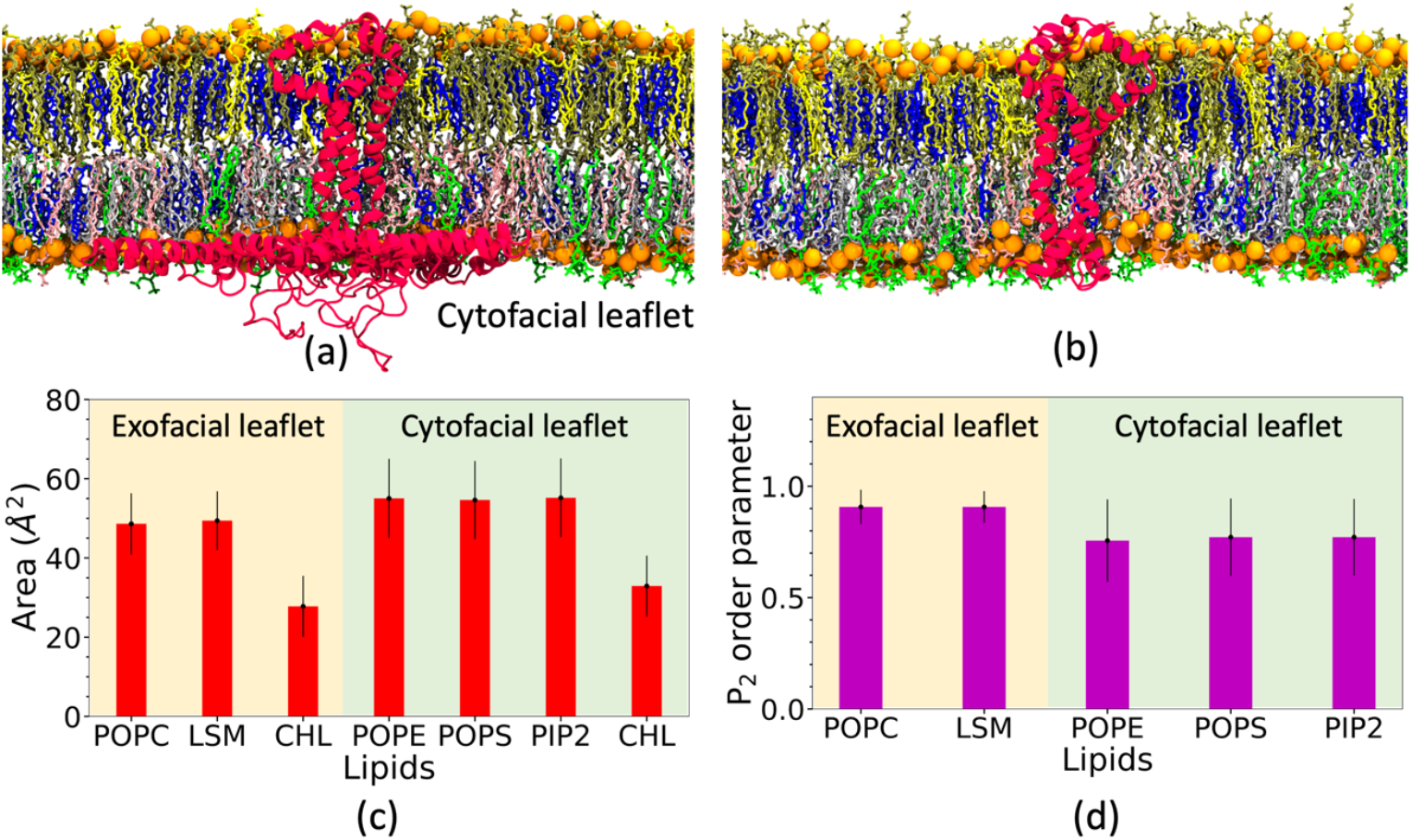
(a) Pictorial representation of the MPER-TMD-CT and (b) MPER-TMD trimers in an asymmetric membrane containing POPC (yellow), LSM (tan), and cholesterol (blue) in the exofacial leaflet, and POPE (gray), POPS (pink), PIP2 (green), and cholesterol (blue) in the cytofacial leaflet. (c) Area per lipid and (d) P_2_ order parameter obtained from the all-atom simulation of the asymmetric bilayer.

HIV-1 entry into the host cell is initiated when gp120s bind to the host cell receptor CD4 and co-receptors CCR5 or CXCR4, triggering a cascade of structural reorganizations of gp41. First, the N-terminal fusion peptide (FP) inserts into the host cell membrane while gp41 remains anchored to the viral membrane through its C-terminal domain, forming a ‘pre-hairpin’ conformation.^12, 13^ Subsequently, the extended ‘pre-hairpin’ conformation folds back to create a ‘hairpin’ structure, where the C-heptad repeat (CHR) domain of the gp41 binds in an antiparallel manner to the trimeric N-heptad repeat (NHR) coiled-coil.^14^ This interaction results in the formation of a stable six-helix bundle that brings the host cell and the viral membranes into proximity.^15^ Then the FP and TM domains of the gp41 disorder both the host and viral membranes, generating a highly curved membrane intermediate that eventually forms a single membrane bilayer.^16^

While the role of the gp41 ectodomain in membrane fusion is well characterized, the function of its membrane-interacting domain remains poorly understood. Previous experimental studies have shown that the MPER and TM domains play crucial roles in HIV-1 entry into host cells. Deletion or mutation in the MPER domain impairs the fusogenic activity of HIV-1.^17^ Truncation of the TM domain also leads to reduced fusogenicity and infectivity.^18^ Additionally, mutations in the MPER and TM domains, as well as deletion of the CT domain, significantly alter the structure of the Env trimer, resulting in changes in binding pattern to broadly neutralizing antibodies (bNAbs).^19-21^ However, truncation or deletion of the CT domain has been reported to have little effect on HIV-1 fusogenicity.^19^

Previous high-resolution structural studies using cryo-EM have characterized the ectodomain of the Env trimer.^22, 23^ However, in those studies the membrane-interacting domain of gp41 was not resolved. Studies using NMR and EPR have determined the structures of the MPER, TM, and CT domains of gp41. The CT domain consisting of amphipathic helices, was found to form a baseplate around the TMD trimer.^11^ However, reported structures of the MPER-TMD region are different based on the conformational topology and oligomeric states (**Figure 1**). Hong and coworkers reported that the MPER-TMD forms a trimeric helix-turn-helix structure in a lipid bilayer.^24^ Chou and coworkers also showed that the MPER-TMD adopts a helix-turn-helix structure and identified a kink near the C-terminal region of the TMD.^20^ In contrast, Bax and coworkers showed that the MPER-TMD forms a monomeric, uninterrupted α-helical structure.^25^ The diverse conformational states observed in previous studies may be ascribed to the variations in protein sequences used, or they might also arise due to the inherent conformational flexibility of the membrane-interacting domain of gp41.

Molecular dynamics (MD) simulations have been widely employed to characterize the structures of proteins in complex membrane environments. Long-timescale MD simulations can effectively sample protein-protein and protein-lipid interactions to predict thermodynamically relevant protein conformational ensembles, whereas experimental studies may only resolve a subset of possible functional forms of a protein complex. In this study, we performed all-atom MD simulations of trimeric complexes of MPER-TMD-CT and MPER-TMD regions of gp41 embedded in a HIV-1 mimetic asymmetric lipid bilayer. Our simulations reveal that the membrane-interacting domain of gp41 adopts diverse conformations depending on the length of the protein construct. We then applied a machine learning based state predictive information bottleneck (SPIB) protocol to investigate conformational ensembles of MPER-TMD trimer. Additionally, we characterized the influence of the CT domain in modulating the lateral organization of the lipids. This study provides an underlying explanation for the diverse structural ensembles reported in previous experimental studies of C-terminal domains of gp41 and elucidates the role of the CT domain in Env recruitment and incorporation at HIV-1 assembly sites.

## Results and discussion

### Simulation of the MPER-TMD-CT and MPER-TMD trimers embedded in an asymmetric lipid bilayer

We simulated an asymmetric bilayer mimicking the composition of the HIV-1 virion using the all-atom CHARMM36m force field. The exofacial leaflet of the asymmetric bilayer is composed of 20 mol% POPC, 40 mol% LSM, and 40 mol% cholesterol, and the cytofacial leaflet consists of 30 mol% POPS, 40 mol% POPE, 15 mol% PIP2, and 15 mol% cholesterol^28, 29^. We used a higher concentration of PIP2 in our simulation compared to the previous HIV-1 lipidomics study to better capture the interactions between PIP2 and other membrane components. To characterize the lateral distribution of lipids across two membrane leaflets, we calculated the liquid crystal order parameter (P_2_) and the area per lipid. The results are shown in **Figure 1**. The P_2_ value ranges from −0.5 to 1, representing lipid tail orientations perpendicular and parallel to the membrane normal, respectively. Previous all-atom simulations have reported P_2_ values above 0.9 for lipids in the liquid-ordered (L_o_) phase and below 0.75 for lipids in the liquid-disordered (L_d_) phase^30, 31^. Compared to previous studies, the high P_2_ value of lipids in the exofacial leaflet indicates the formation of a L_o_ phase, while the lower P_2_ value in the cytofacial leaflet suggests the formation of a L_d_ phase. We also calculated the area per lipid using Voronoi tessellation based on the positions of individual lipid tails (**Figure 1**). The smaller area per lipid observed in the exofacial leaflet compared to the cytofacial leaflet reflects greater lipid tail order in the exofacial leaflet and highlights the cholesterol condensation effect. We also computed the spatial distribution of the P_2_ order parameter in a plane parallel to the membrane surface. The results indicate no phase separation of the lipids in either leaflet (**Supplementary Figure 1**).

Upon establishing the HIV-1 membrane properties, we simulated the membrane-interacting domains of gp41 by embedding them in the asymmetric bilayer of the same composition. The MPER of the gp41 was placed in the exofacial leaflet, and the CT domain was placed in the cytofacial side of the asymmetric bilayer. Two protein constructs of gp41: residues 660 to 716 corresponding to the MPER-TM domain, and residues 660 to 856 corresponding to the MPER-TMD-CT domain, were studied. For each protein construct, three independent simulations were performed. We calculated the insertion depth of the protein in the membrane (**Supplementary Figure 2**). The CT domain of gp41 was found to be stable at the interface of cytofacial leaflet and water. The MPER domain predominantly interacts with the water on the exofacial side of the membrane, as observed in previous experimental studies.^11, 24^ However, in the absence of the CT domain, the MPER domain was found to be more exposed to water.

Chou and coworkers measured the backbone dynamics of the MPER-TM domain of gp41 using NMR relaxation rates and reported higher dynamics in the MPER and the C-terminal region of the TMD.^32^ We calculated the root-mean-square fluctuation (RMSF) of the protein to analyze the dynamics of residues in different protein domains (**Supplementary Figure 2**). The results show higher fluctuation in the MPER and the C-terminal region of TMD, consistent with the previous observations. Higher dynamics were also observed in the loop region of the CT baseplate.

### PIP2 lipids preferentially bind to the basic residues in the CT domain

The CT domain consists of amphipathic helices that wrap around the C-terminal end of the TMD trimer (**Supplementary Figure 3**). The CT domain plays a crucial role in Env trafficking and clustering at the HIV-1 assembly sites.^33, 34^ Roy *et al*. showed that the Gag assembly induces the aggregation of the Env, and deletion of the CT domain abrogates Gag’s influence on Env recruitment.^34^ Kräusslich and coworkers demonstrated that the Env aggregation process depends on both the matrix (MA) domain of Gag and the CT domain of gp41.^35^ Envs were found to aggregate near the periphery of the Gag assembly site. They also showed that the Env does not form clusters in the absence of Gag, and deletion of the CT domain leads to a scattered distribution of Env. Additionally, depletion of PIP2 lipids causes disintegration of the Gag lattice, resulting in the scattering of Env clusters.^33^ Based on these observations, they hypothesized that an indirect interaction between the MA domain of Gag and the CT domain of gp41 via the formation of a PIP2-rich membrane microdomain leads to Env recruitment at the HIV-1 assembly site.

To characterize the interaction between lipids and gp41 on the cytofacial side of the membrane, we calculated the density distribution of the CT domain and PIP2 lipids on the *xy*-plane parallel to the membrane surface. The results are shown in **Figure 3**. The TMD trimer was centered on the *xy*-plane, and a vector defining the center of mass (COM) of the TMD trimer to the COM of TM helix A was aligned to the positive *x*-axis. The CT domain baseplate was found to be stable around the TM helices (**Figure 3b**). A higher local density of PIP2 was observed around the CT domain baseplate (**Figure 3c**). We also calculated the contact fraction between the CT domain and PIP2 lipids (**Figure 3d**). The resulting binding pattern shows that the interactions between the CT domain and PIP2 lipids are mediated by basic residues in the CT domain. This finding supports the hypothesis of a lipid-mediated mechanism of Env clustering at the HIV-1 assembly site, specifically through the formation of a PIP2-rich membrane microdomain.

**Figure 3:**
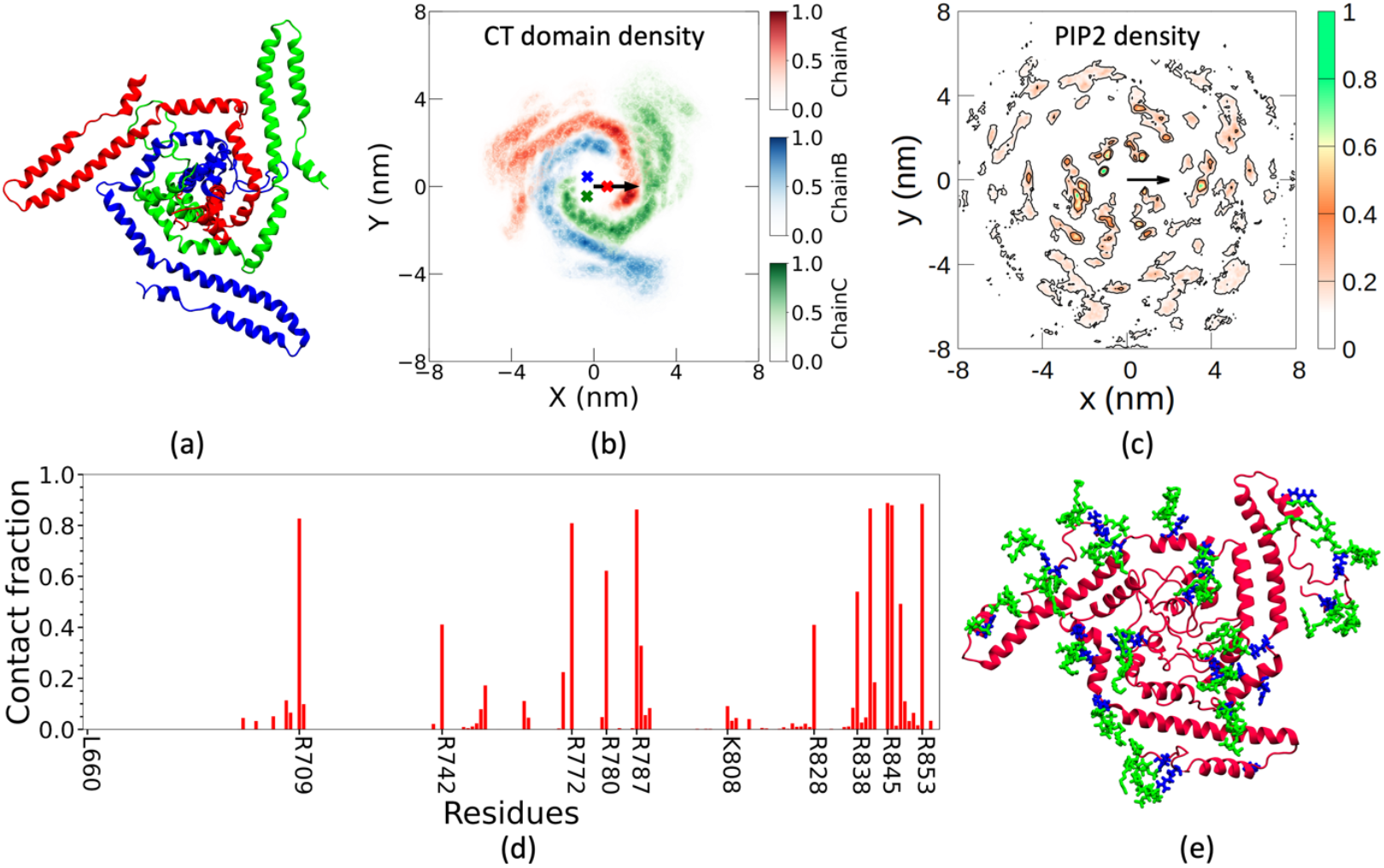
(a) Schematic representation of the three chains of the MPER-TMD-CT domain of gp41. (b) Probability density of the CT domains in the *xy*-plane. The center of mass (COM) of the TM helices are shown as dots. (c) Probability density of PIP2 in the cytofacial leaflet around the protein. The COM of the three TM domains of the trimer was centered on the *xy*-plane. The arrow represents the vector defined by the COM of the three TM helices through the COM of the TM helix A. (d) Interactions between the CT domains and PIP2, where a contact fraction value of 1 indicates a contact maintained by a residue and a PIP2 lipid throughout the simulation trajectory. (e) Pictorial representation of the PIP2 lipid interactions with the CT domain. Protein, basic residues, and PIP2 are represented in red, blue, and green, respectively.

### GXXXG motifs are crucial in mediating interactions between transmembrane helices

Transmembrane domain anchors the Env to the membrane and is the sole mediator of physical coupling between the CT domain and the ectodomain of the Env. The TMD also plays a crucial role in maintaining the structure and function of the Env. Hunter and coworkers showed that truncation of the TM domain of gp41 abrogates the fusogenic properties of Env.^18^ Chou and coworkers demonstrated that mutation of the G690XXXG694 motif in the TM helix and deletion of the C-terminal region of the TMD completely disrupt the trimeric structure of gp41, resulting in a monomeric form.^20^ However, NMR studies observed no significant changes in interactions between the TM domains of protein constructs with or without the CT domain.^32^

We characterized the interaction between the TM helices in MPER-TMD and MPER-TMD-CT trimers. In a simulation of TM protein in a membrane bilayer, the movement of the TM helices can be described as motion in the *xy*-plane parallel to the bilayer surface. The probability density of the TM helices obtained from simulations is shown in **Figure 4**. One of the helices in the trimer was centered on the *xy*-plane, and the COM690-to-Gly690 vector of the centered helix was aligned to the positive *x*-axis, where COM690 is defined as the COM of residues 688 to 692. The probability density of the two other helices relative to the centered helix is shown in **Figure 4**. The density observed along the positive *x*-axis indicates that the interaction between the TM helices is mediated by the Gly690XXXGly694 motif. These results are in line with previous experimental mutagenesis study of gp41.^20^ The contact map defining the interactions between the TM domains of the gp41 trimer is shown in **Supplementary Figure 4**. The results show that in addition to the Gly690XXXGly694 motif, the Ilu693XXXIlu697 motif also plays an important role in stabilizing the gp41 trimer. Additionally, as observed in a previous NMR study, the interaction pattern between the TM helices is not affected by the presence of the CT domain.^32^

**Figure 4:**
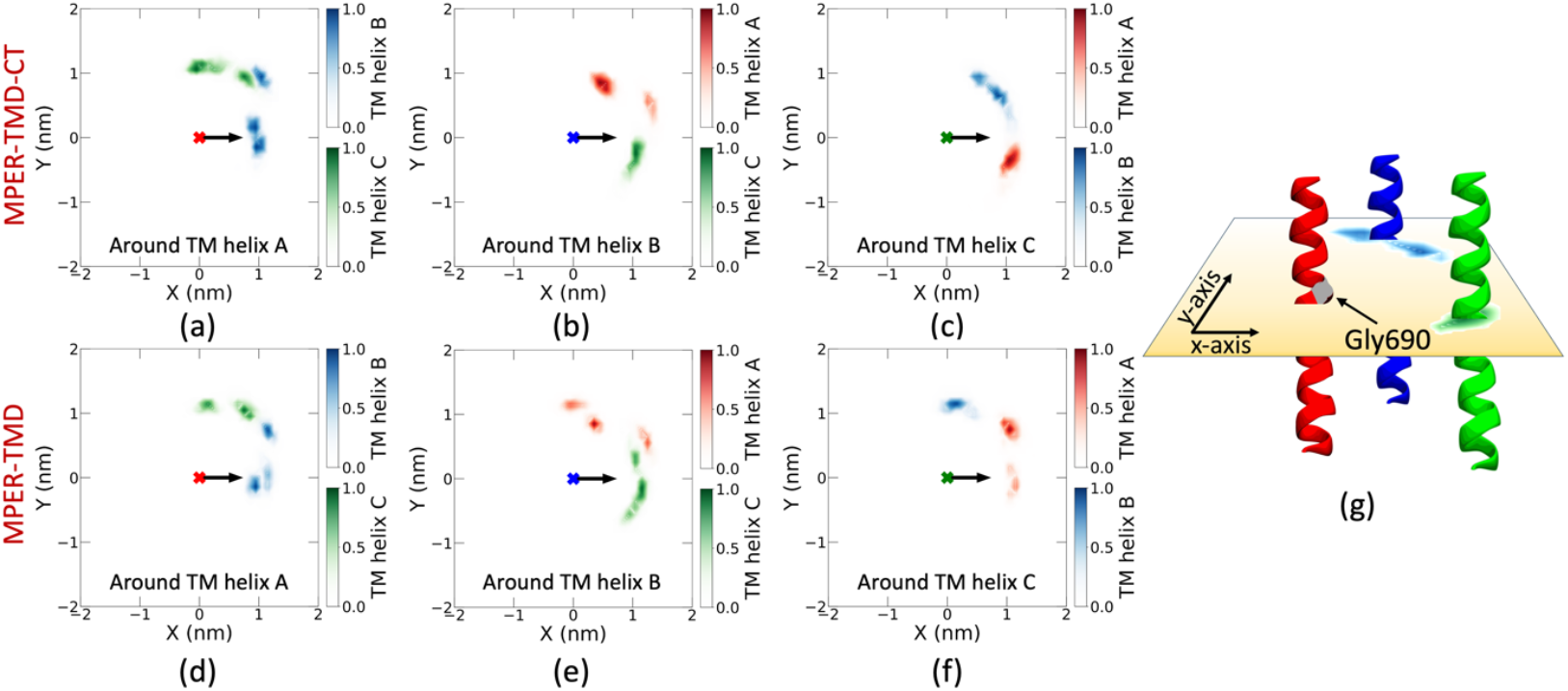
Interaction between three transmembrane helices, where one helix was centered in the *xy*-plane and the position of the other two helices were shown as a colormap. Probability density of (a, d) TM helices B and C around helix A, (b, e) TM helices C and A around helix B, and (c, f) TM helices A and B around helix C obtained from the simulations of MPER-TMD-CT and MPER-TMD trimers, respectively. The arrow represents the COM^690^-to-Gly690 vector of the centered helix. The COM^690^ of a helix is defined as the center of mass of residues 688 to 692. (g) Schematic representation of the interactions between three TM helices.

### Diverse conformations of MPER-TMD were observed in both simulations and experiments

Several experimental studies have characterized the structure of the membrane-interacting domain of gp41 and reported diverse conformational states of the MPER-TMD. Bax and coworkers studied residues 677-716 of gp41 in DMPC/DHPC bicelles and reported a monomeric α-helical conformation of the protein construct.^25^ Hong and coworkers characterized the structure of residues 665 to 704 in a bilayer containing POPC, POPE, POPS, SM, and cholesterol and reported that the MPER-TMD adopts a trimeric helix-turn-helix conformation, with a hinge located at the junction of the MPER and TMD.^24^ Reinherz and coworkers studied the structure of the MPER-TMD in DPC micelle and reported that residues 664-672 of the MPER domain form an α-helix, connected to the α-helical TMD (residue 675-683) via a short hinge.^26^ Nieva and coworkers studied MPER-TMD in hexafluoroisopropanol and reported that the MPER and the N-terminal region of TMD, and the C-terminal region of the TMD both form an α-helical structure. However, a helical kink was observed near the G690XXXG694 motif of the TMD.^27^ Chou and coworkers characterized the complete membrane-interacting region of gp41 in DMPC/DHPC bicelles in a series of studies.^11, 20, 32^ They reported that the MPER-TMD adopts a helix-turn-helix conformation, with a hinge at the MPER and TMD junction, and also identified a helical kink at residue 703 near the C-terminus of the TMD. These diversities in the reported structure of the MPER-TMD region of gp41 highlight the conformational flexibility of the domain. In an experimental study, Chen and coworkers also showed that deletion of the CT domain causes significant structural changes in Env, which in turn affect the binding affinities of various bNAbs to Env.^19^

We characterized the conformational ensemble of the MPER-TM domain of gp41 observed in simulations with and without the CT domain. First, we analyzed the secondary structure of the residues within the MPER-TMD region (**Supplementary Figures 5 and 6**). The results indicate that both the MPER and TM domains form stable α-helical structures. However, two disordered regions were observed, one at the junction of MPER and TMD, and another at the C-terminal region of TMD near residue 700. The disordered residues lead to the formation of hinges between MPER and the N-terminus of the TMD (ϕ_top_) and between the N-terminal and C-terminal regions of TMD (ϕ_bottom_). The distributions of the top hinge angle (ϕ_top_) and bottom hinge angle (ϕ_bottom_) illustrate the diversity of MPER-TMD conformations (**Figure 5**). The results indicate that the CT domain restricts the bending of the C-terminal domain of the TMD, leading to a higher ϕ_bottom_ value. In the absence of the CT domain, the C-terminal of the TMD favors bending, resulting in a smaller ϕ_bottom_ value and the formation of an unhinged MPER-TMD conformation. These changes in the top and bottom hinge angle patterns also help relieve the hydrophobic mismatch of the transmembrane domain. Additionally, we computed the crossing angle distribution between the TM helices (**Figure 5**). However, in contrast to a previous hypothesis by Chou and coworkers,^32^ we did not observe significant differences in the crossing angle distribution of protein structures with and without the CT domain.

**Figure 5:**
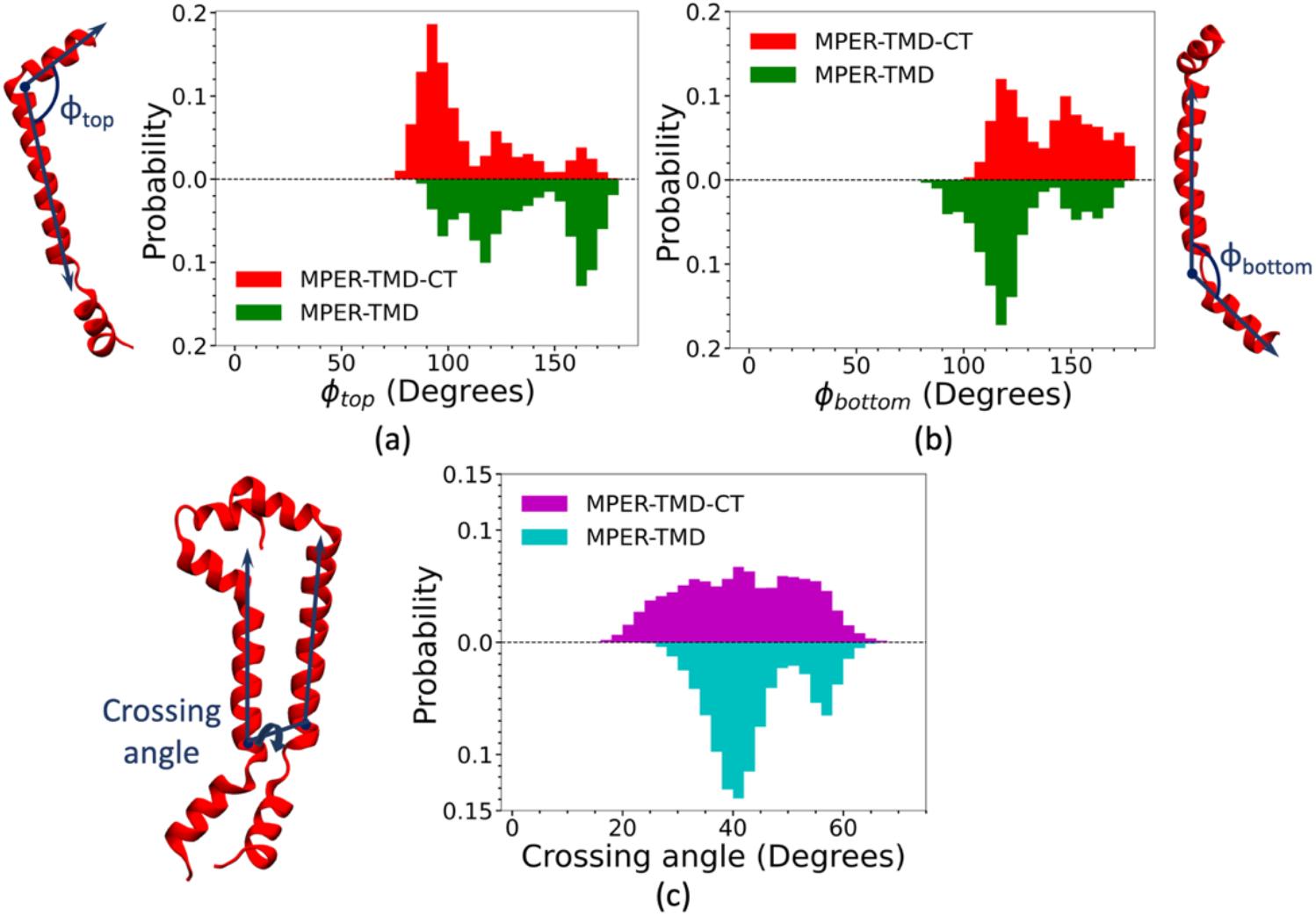
Probability density of the (a) top hinge angle (ϕ_top_), (b) bottom hinge angle (ϕ_bottom_), and (c) crossing angle between TM domains, characterizing the structural ensembles of MPER-TMD-CT and MPER-TMD trimers. Diverse protein conformations were observed in the simulations.

### Characterizing MPER-TMD configurations using an artificial intelligence based state predictive information bottleneck (SPIB) protocol

The hinge angle distributions provide insight into the structures of the MPER-TMD monomer. However, the structural characterization of MPER-TMD trimers remained elusive. We therefore applied the AI-based SPIB protocol^36^ to analyze the conformational ensemble of MPER-TMD trimers. This protocol was used to derive a two-dimensional collective variable (CV) describing the structures of the trimer. Configurations of the MPER-TMD domain obtained from the unbiased simulations were used to train a neural network (NN) model. A total of 36 descriptors were used to define each MPER-TMD trimer structure (**Supplementary Figure 7**). The SPIB protocol uses *Δt* as a hyperparameter to incorporate state information at time *t* in predicting the state information at time *t*+ *Δt*. We utilized data from six independent unbiased simulations of MPER-TMD-CT and MPER-TMD trimers to extract the descriptor values and initial state labels over time. The initial state label for each trimer was assigned based on the structure of its constituent MPER-TMD monomers. Based on the distribution of hinge angles, the diverse structural ensembles of MPER-TMD monomer were categorized into 4 distinct states (**Figure 6a**). Using this classification, 20 initial states were assigned to describe MPER-TMD trimer structures. For example, a state 1-1-2 refers to a trimer structure where two monomers represent state 1, and the third monomer represents state 2. The encoder and decoder NN models were then trained using the initial state labels, which are iteratively refined to yield a converged estimation of the final states. The projection of the initial data points onto the SPIB CVs is shown in **Figure 6b**. The results show five important states that represent the complete structural ensembles of the MPER-TMD trimer.

**Figure 6:**
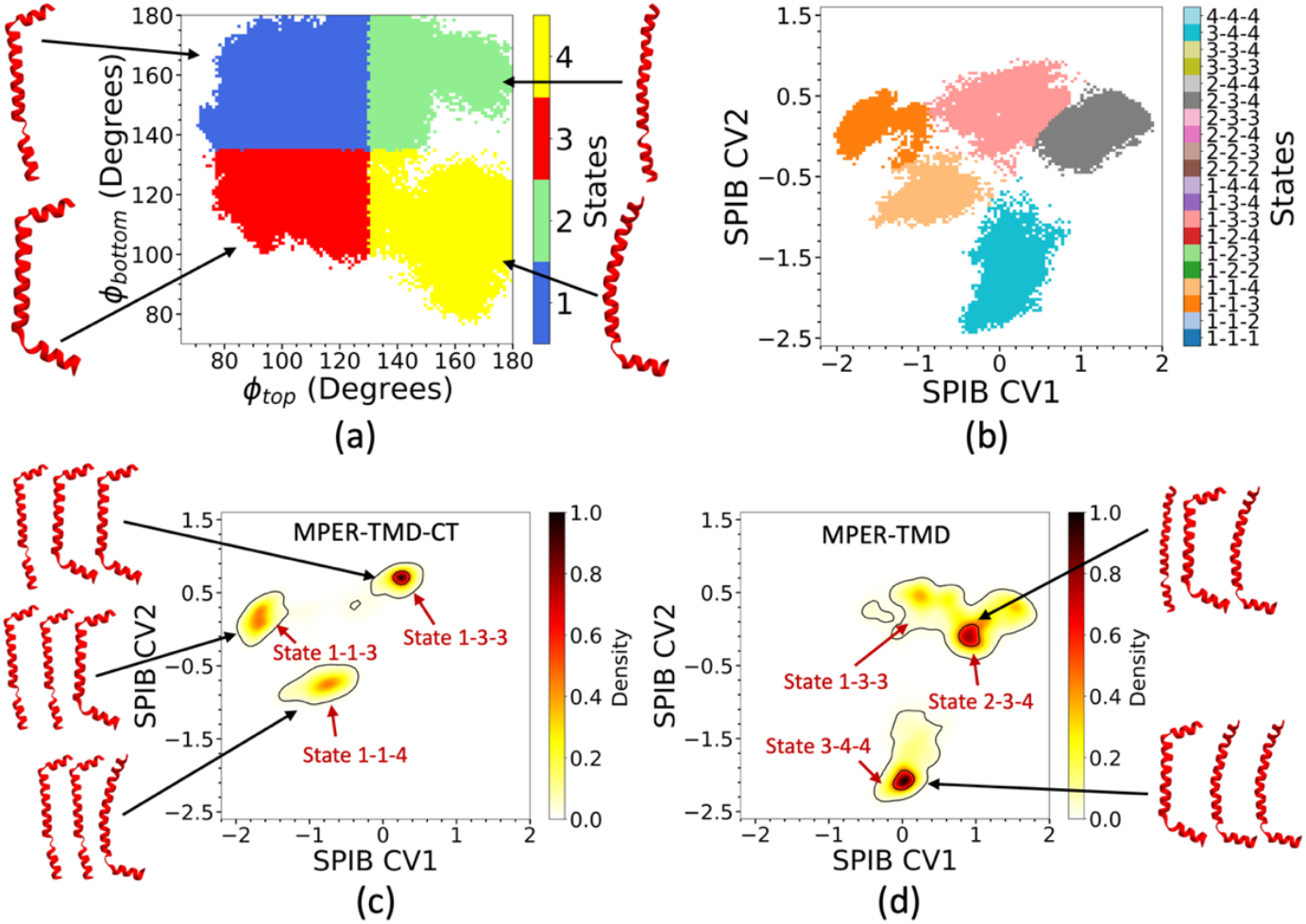
(a) Four initial states were defined based on the value of top (ϕ_top_) and bottom (ϕ_bottom_) hinge angles of MPER-TMD monomer. A total of 20 initial states were defined to identify a conformation of MPER-TMD trimer. (b) Converged state representations of the MPER-TMD trimer conformations onto the SPIB CVs. Probability densities as a function of the SPIB CVs characterizing the timer conformations of the MPER-TMD domain obtained from simulations of the (c) MPER-TMD-CT and (d) MPER-TMD trimers.

To better understand the influence of the CT domain in modulating the structure of the MPER-TMD domain, we calculated the probability density of the MPER-TMD-CT and MPER-TMD trimers onto the two-dimensional SPIB CV (**Figures 6c and d**). The results show that the CT domain significantly modulates the structure of the MPER-TM trimer. In the presence of the CT domain, the MPER-TMD predominantly adopts structures corresponding to states 1-3-3, 1-1-3, and 1-1-4. Whereas, the MPER-TMD trimer predominantly forms states 1-3-3, 2-3-4, and 3-4-4 in the absence of the CT domain. The crossing angle distributions of different trimer states are shown in **Supplementary Figure 8**.

By adding important additional understanding to the previous experimental MPER-TMD trimer structures, where all monomers were reported to adopt the same conformation,^11, 24^ our results demonstrate that the MPER-TMD monomers can adopt diverse conformational states while forming thermodynamically stable trimer configurations.

In comparison with previous experimental studies, the structure of the MPER-TM domain reported by Hong and coworkers,^24^ as well as by Reinharz and coworkers,^26^ resembles the monomer state 1 used in this study. The structures reported by Bax and coworkers^25^ and Chou and coworkers^11, 32^ correspond to monomer states 2 and 3, respectively. A comparison between prior studies and the present work indicates that all reported conformations above are relevant for defining the complete structural ensemble of the MPER-TMD. However, we did not observe an α-helical conformation of MPER-TMD with a kink near the G690XXXG694 motif, as reported by Nieva and coworkers.^27^ We also note that the MPER-TM domain of gp41 forms a stable trimeric structure in a membrane bilayer, consistent with the findings of Hong and coworkers and Chou and coworkers. Taken together, this study offers an explanation for the diverse conformational states of the membrane-interacting domain of gp41 observed in previous studies. Due to the conformational flexibility of the MPER-TMD, it can adopt multiple distinct configurations, and prior experimental studies captured only a subset of the many thermodynamically stable conformations.

## Conclusions

In this study, we characterized the structure and functions of the membrane-interacting domain of gp41 which helps to understand the mechanism of Env trafficking. Using all-atom MD simulation, we modeled an asymmetric membrane mimicking the HIV-1 membrane composition. The exofacial leaflet of the asymmetric bilayer was observed to form a liquid-ordered phase, and the cytofacial leaflet was observed to form a liquid-disordered phase at room temperature. Subsequently, we simulated the structure and dynamics of MPER-TMD and MPER-TMD-CT domains of gp41 in the same model HIV-1 membrane. The MPER was observed to localize at the exofacial leaflet-water interface, whereas the CT domain forms a stable baseplate around the TM domain at the cytofacial leaflet-water interface. The transmembrane domains of gp41 were observed to form a stable trimeric structure both with or without the CT domain, with the interactions between the TM helices mediated by the G690XXXG694 and Ilu693XXXIlu697 motifs, in agreement with previous studies.^11, 20^

Earlier studies have shown that Env trafficking and incorporation at the HIV-1 assembly site depend on both the CT domain of gp41 and the MA domain of Gag, with Gag forming clusters on the cytofacial leaflet in a PIP2-dependent manner.^33, 35^ To characterize the interaction between lipids and the CT domain, we analyzed the lateral distribution of lipids in the cytofacial leaflet around the protein. We observed lipid demixing and the accumulation of PIP2 lipids around the CT baseplate and found that the basic residues in the CT domain are crucial in facilitating interactions between PIP2 lipids and gp41. These results support the hypothesis of Env trafficking near the Gag assembly site via the formation of a PIP2-rich membrane microdomain.

Collective variables were derived using the AI-based machine learning SPIB protocol to analyze the structures of the MPER-TMD trimer. Data obtained from extensive unbiased simulations was used to train the deep NN models. Projection of the probability density of the MPER-TMD trimers onto the SPIB-derived CVs reveals that each monomer can adopt different conformational states while forming a stable trimeric structure. The CT domain was found to modulate the overall structure by restricting the bending of the C-terminal domain of the TMD, which also leads to the formation of a hinged MPER-TMD conformation. Assessment of the MPER-TMD structures from the simulations shows good agreement with previous experimental studies.^11, 20, 24-26^ The membrane-interacting domain of gp41 can adopt diverse structures due to its inherent conformational flexibility. Apparent discrepancies with earlier experimental studies arose because they characterized one of the many functional forms of MPER-TMD. In this study, we have provided a more complete characterization of the thermodynamically relevant structures of the membrane-interacting domain of gp41.

It was previously conjectured that the highly conserved MPER and CT domains are critical in modulating the antigenic structure of the Env.^19, 21^ The comprehensive understanding of the different conformational states of the MPER-TMD-CT domain of gp41 obtained from this study may enable the rational design of therapeutic compounds to manipulate the structure of Env for effective antibody responses.

## Methods

### All-atom simulation of protein-embedded lipid bilayers

We simulated an asymmetric membrane bilayer composed of six different lipids. The exofacial leaflet consisted of 20 mol% POPC, 40 mol% LSM, and 40 mol% cholesterol, while the cytofacial leaflet consisted of 40 mol% POPE, 30 mol% POPS, 15 mol% PIP2, and 15 mol% cholesterol. We also simulated the trimeric MPER-TMD-CT and MPER-TMD regions of gp41 by embedding them into the asymmetric bilayer of the same composition. The structures of the membrane-interacting domain of gp41 were obtained from PDB ID 7LOI.^11^ The initial configurations of the protein-embedded membrane bilayers were prepared using the CHARMM-GUI membrane builder.^37^ Membrane bilayers without protein and with protein were solvated using 50 and 100 water molecules per lipid, respectively, defined by the TIP3P water model.^38^ A KCl concentration of 0.15 M was used. The protein-embedded membrane bilayers were simulated using the CHARMM36m force field.^39^ All protein-membrane systems studied are listed in **Supplementary Table 1**. Each bilayer containing protein was equilibrated for 100 ns following the CHARMM-GUI protocol.^40^ Production runs were performed in the NPT ensemble using a leap-frog integrator with a 2 fs timestep. Three independent simulations were performed for each protein-embedded membrane bilayer. Membranes containing MPER-TMD-CT and MPER-TMD trimers were simulated for a total of 6 μs and 9 μs, respectively. The Lennard-Jones potential was calculated using a cutoff of 1.2 nm with a force-switch function between 1.0 and 1.2 nm. The electrostatic potential was calculated using the particle mesh Ewald method. The temperature of the simulations was maintained at 303 K using the Nose-Hoover thermostat. The pressure during equilibration and production runs was maintained at 1 bar using the semi-isotropic Berendsen and Parrinello-Rahman barostats. All simulations were carried out using the GROMACS software.^41^

The liquid crystal order parameter (P_2_) of the lipids was calculated using the angle (θ) between the lipid director vector and the membrane normal.

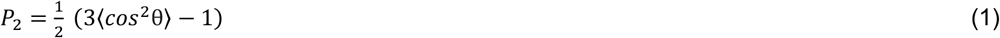

For POPC, POPE, POPS, and PIP2, the director vectors were defined by the C1 through C16 atoms and C1 through C18 atoms. For LSM, the director vectors were defined by the C1 through C16 atoms and C1 through C18 atoms.

### Developing collective variables using the SPIB protocol

The AI-based SPIB protocol^36^ was used to characterize the different conformational ensembles of the MPER-TMD trimer of gp41. SPIB employs an informational bottleneck approach that uses input descriptors *D* to seek a concise representation *z* that retains maximum information about the target *s*. A model using the SPIB protocol can be trained with initial state labels, which are iteratively refined to obtain the final state labels.

Descriptors defining the configurations of the trimer obtained from the unbiased simulations are denoted as *D*_*1*_, *D*_*2*_, *D*_*3*_, *…*., *D*_*M+*x_, and the initial state labels are denoted as *s*_*1*_, *s*_*2*_, *s*_*3*_, *…*., *s*_*M+x*_, where *D*_*t*_ and *D*_*t+x*_ represent the values of descriptors and *s*_*t*_ and *s*_*t+x*_ represent the initial state labels at an arbitrary time *t* and *t*+*Δt*, respectively. The *Δt* is the time delay. The SPIB model can be trained using data obtained from a large unbiased trajectory by maximizing the objective function

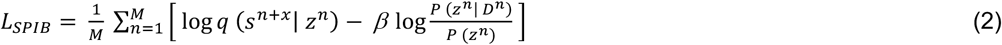

where *M* is the total number of data points. The first term of the objective function defines the capability of the model to predict the desired final state label, while the second term acts as a regularizer of information from the input descriptors *D* to *z*. The parameter *β* controls the trade-off between these two terms. The terms *q* (*s*^*n*+*x*^| *z*^*n*^), *P* (*z*^*n*^| *D*^*n*^), and *P* (*z*^*n*^) are calculated using the deep neural network (NN) models.

Each MPER-TMD trimer configurations were described by 36 descriptors. Six descriptors defined the top hinge angles (ϕ_top_) and bottom hinge angles (ϕ_bottom_) of the three chains. Three descriptors defined the crossing angles between the TM helices A and B, TM helices B and C, and TM helices C and A. Finally, 27 descriptors captured the inter-chain distances between COM_helix1_, COM_helix2_, and COM_helix3_ for each chain of the MPER-TMD trimer (**Supplementary Figure 7**). Protein conformations were saved every 200 ps during the unbiased simulations to generate the dataset for the training of the SPIB model. Two hidden layers with 32 neurons each were used to model both the encoder and decoder. A *β* value was set to 0.001, and a time delay *Δt* was chosen to be 400 ps. The NN model was optimized using the ADAM optimizer with a learning rate of 0.001. The SPIB code used in this study was adapted from the work of Tiwary and coworkers.^36^

## Supporting Information

The supplementary information contains (**Table 1**) Systems simulated in this study; (**Figure 1**) Analysis of the asymmetric bilayer; (**Figure 2**) Insertion depth and RMSF of proteins; (**Figure 3**) Helical wheel representation of the CT domain; (**Figure 4**) Contact map of TM helices; (**Figure 5 and 6**) Secondary structure analysis of MPER-TMD-CT and MPER-TMD; (**Figure 7**) Pictorial representation of descriptors; (**Figure 8**) Crossing angle distribution of different MPER-TMD trimer conformations.

**Figure 7:**
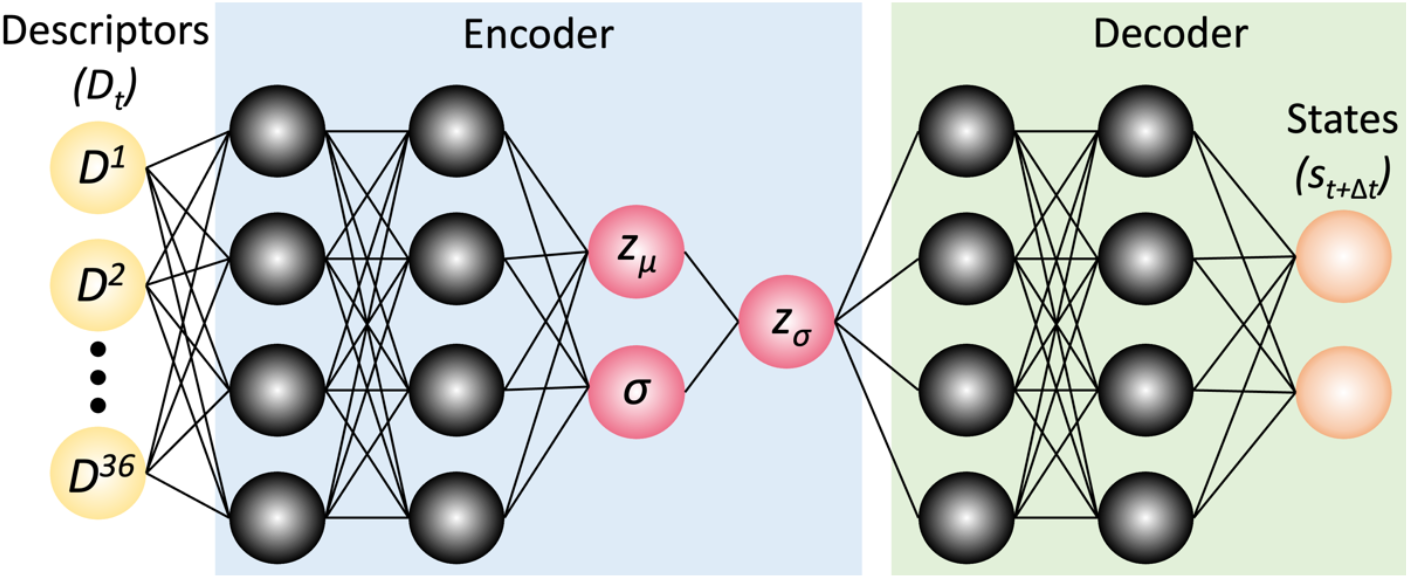
Schematic representation of the AI-based SPIB protocol used in this study. A total of 36 descriptors were used to characterize a MPER-TMD trimer conformation.

## Acknowledgments

This research was supported by the National Institute of Allergy and Infectious Diseases (NIAID) of the National Institutes of Health (NIH) through grant R01AI178850. The content is solely the responsibility of the authors and does not necessarily represent the official views of the National Institutes of Health. The authors gratefully acknowledge computational resources provided by the University of Chicago Research Computing Center and the Frontera supercomputer at the Texas Advanced Computer Center funded by the National Science Foundation (OAC-1818253).

