## Supplementary Information for "Structural Heterogeneity of the Membrane-Interacting Region of the HIV-1 Envelope Glycoprotein"

**Table 1:** Protein-membrane systems simulated in this study.

| System | Number of lipids | Number of replicas | Total simulations (μs) |
|---|---|---|---|
| HIV-1 mimetic asymmetric bilayer | 2271 | 1 | 1 |
| MPER-TMD-CT trimer in a HIV-1 mimetic asymmetric bilayer | 1135 | 3 | 6 |
| MPER-TMD trimer in a HIV-1 mimetic asymmetric bilayer | 454 | 3 | 9 |

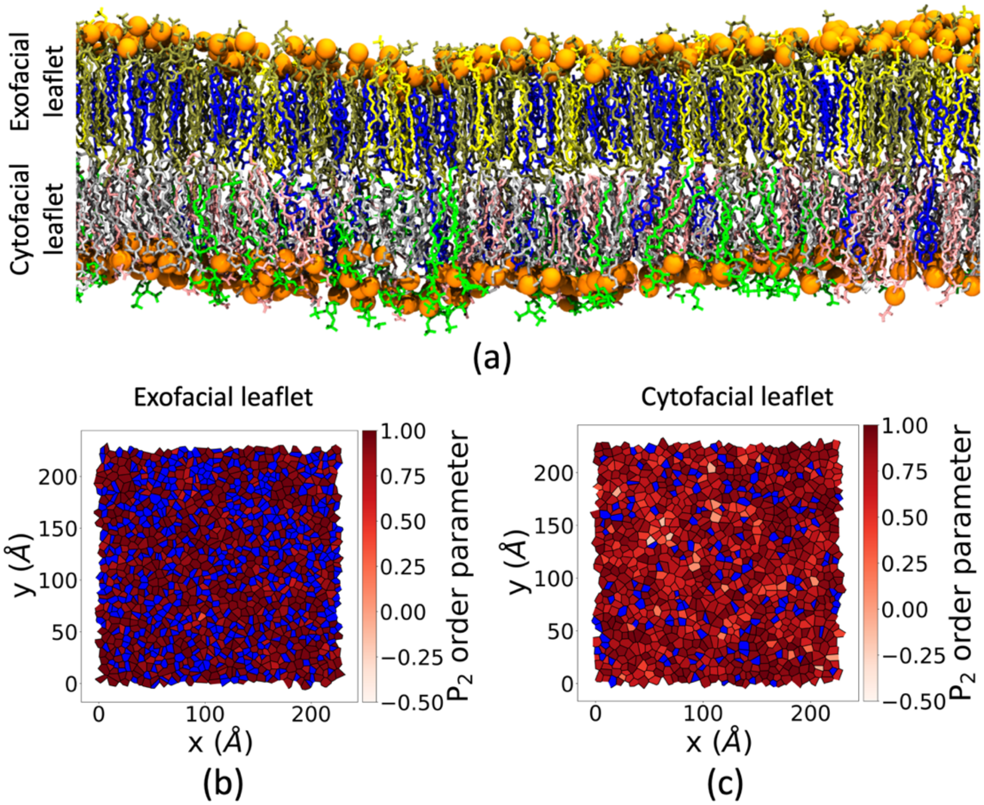


**Supplementary Figure 1:** (a) Schematic representation of the HIV-1 mimetic asymmetric bilayer containing POPC (yellow), LSM (tan), and cholesterol (blue) in the exofacial leaflet, and POPE (gray), POPS (pink), PIP2 (green), and cholesterol (blue) in the cytofacial leaflet. Lateral distribution of the $P_2$ values in the (b) exofacial and (c) cytofacial leaflets of the asymmetric bilayer. No phase separation of lipids was observed in either leaflet of the asymmetric bilayer.

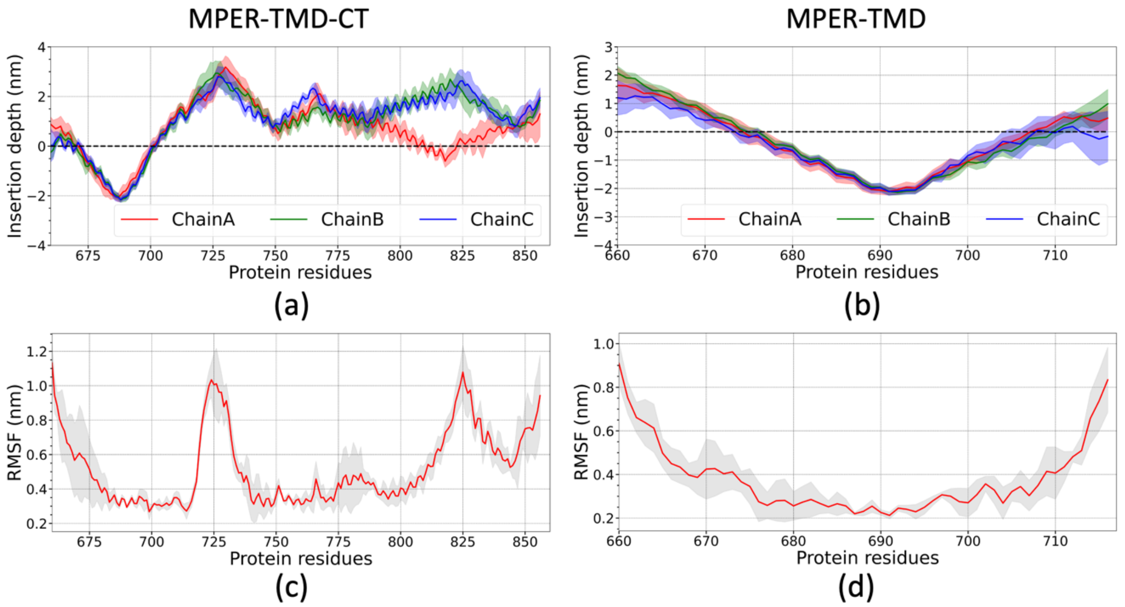


**Supplementary Figure 2:** Average depth of insertion of the (a) MPER-TMD-CT and (b) MPER-TMD trimers in membrane bilayer. Root mean square fluctuation (RMSF) of the (c) MPER-TMD-CT and (d) MPER-TMD region of gp41.

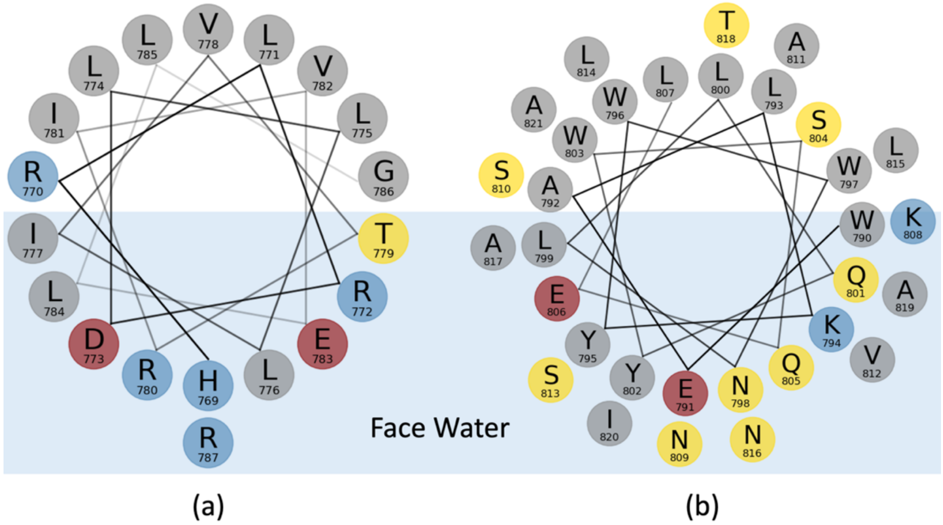


**Supplementary Figure 3:** Helical wheel representations of the α-helices comprising (a) residues 769–787 and (b) 790–821, which form the CT domain of gp41. Basic, acidic, polar, and nonpolar residues are represented by blue, red, gold, and gray colors, respectively. The α-helices in the CT domain are amphipathic in nature.

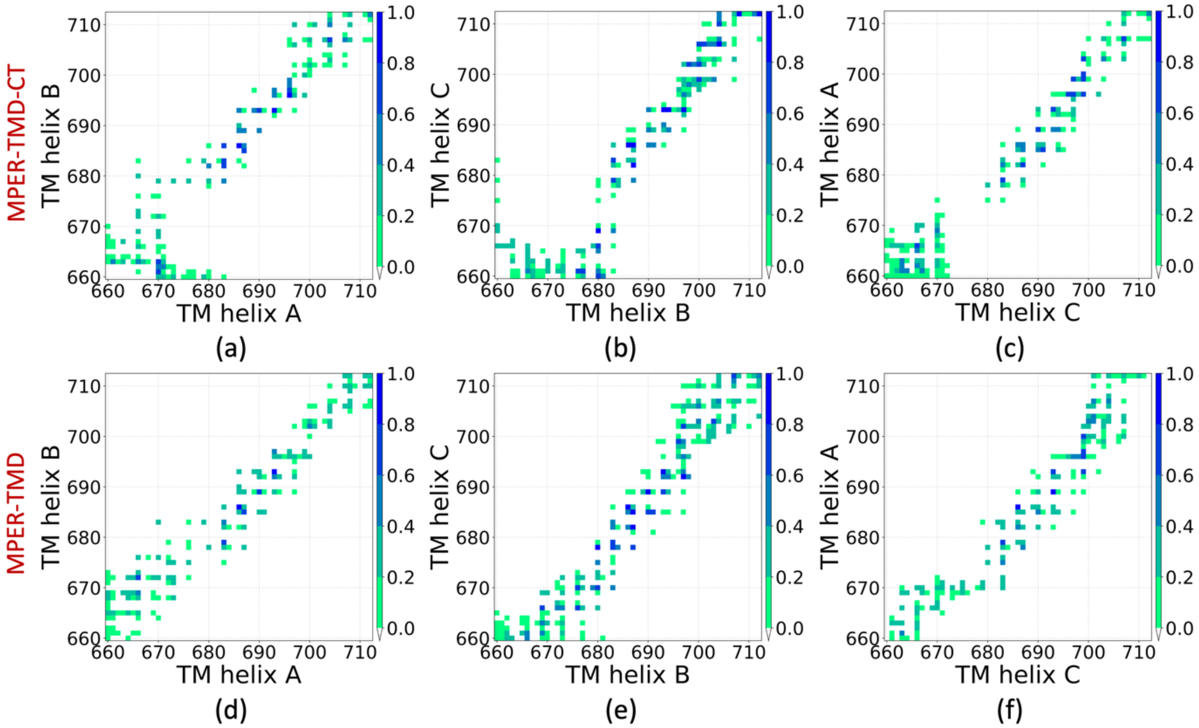


**Supplementary Figure 4:** Contact map defining the interactions between (a, d) TM helix A and B and (b, e) TM helix B and C, and (c, f) TM helix C and A of the MPER-TMD-CT and MPER-TMD trimers. A cutoff distance of 5 Å was used to identify contacts between residues. Similar interactions between TM helices were observed in both MPER-TMD-CT and MPER-TMD trimers.

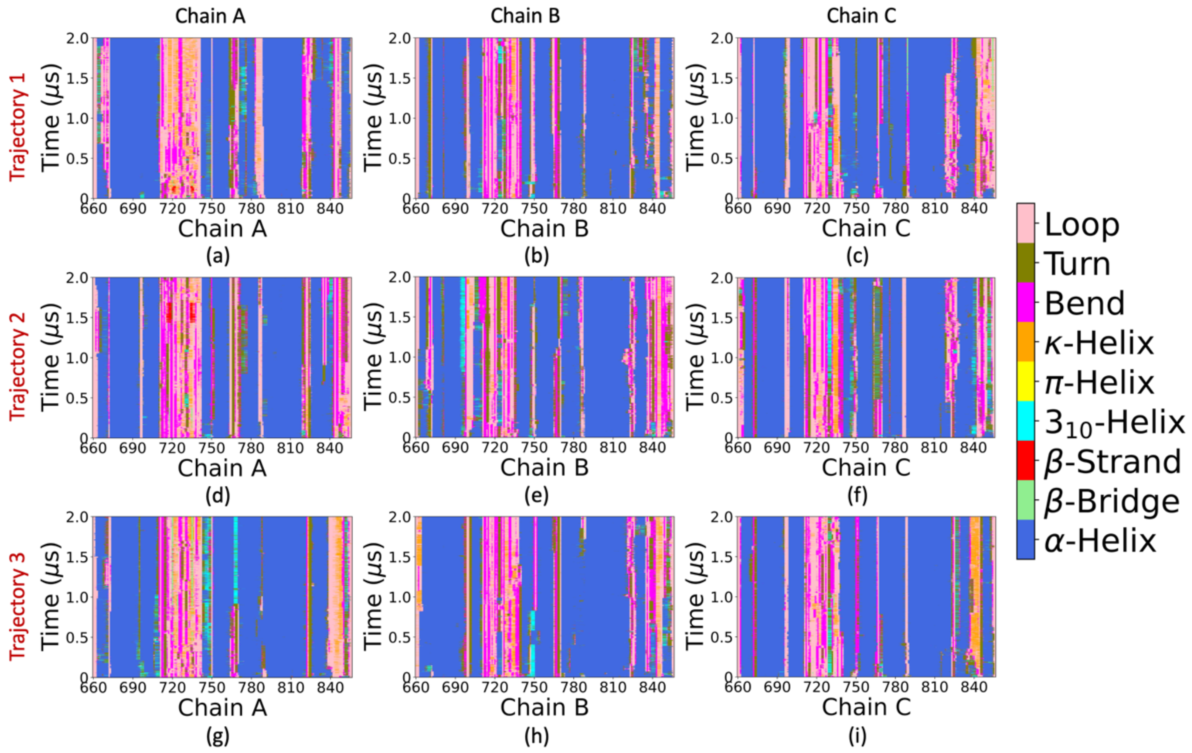


**Supplementary Figure 5:** Secondary structure of the residues in (a, d, g) chain A, (b, e, h) chain B, and (c, f, i) chain C of the MPER-TMD-CT trimer obtained from independent simulation trajectories 1, 2, and 3, respectively.

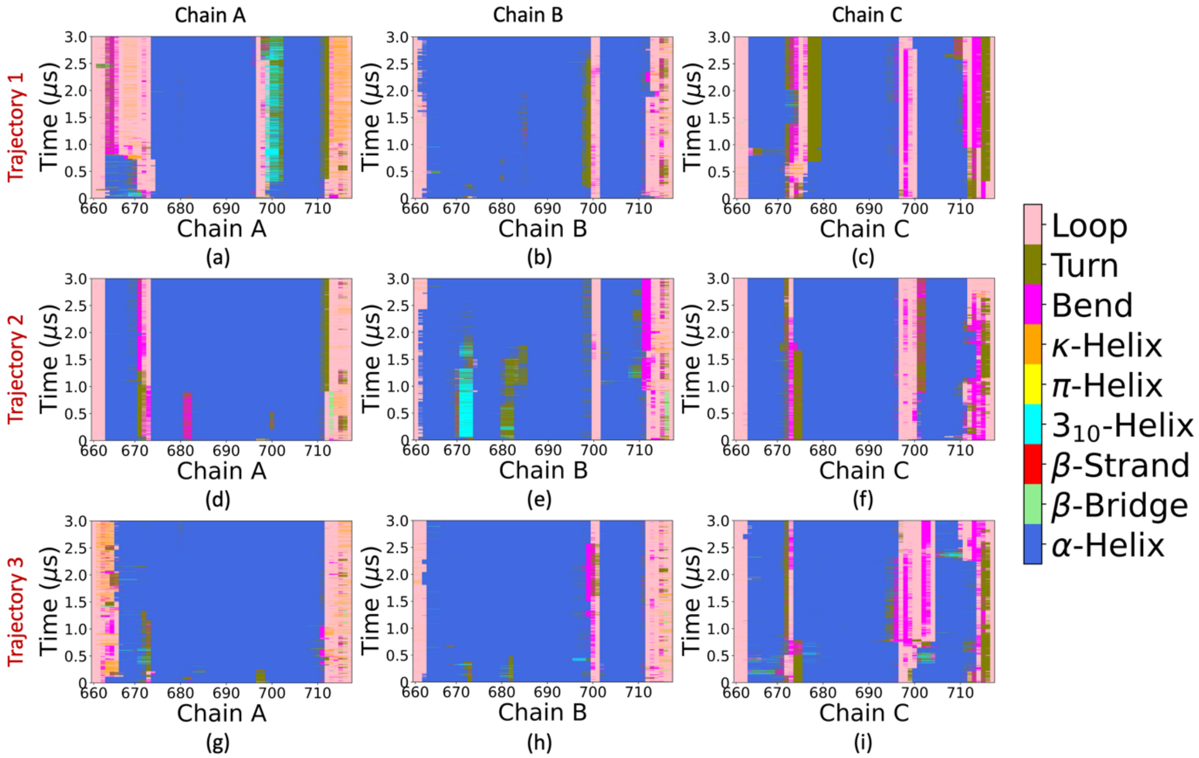


**Supplementary Figure 6:** Secondary structure of the residues in (a, d, g) chain A, (b, e, h) chain B, and (c, f, i) chain C of the MPER-TMD trimer obtained from independent simulation trajectories 1, 2, and 3, respectively.

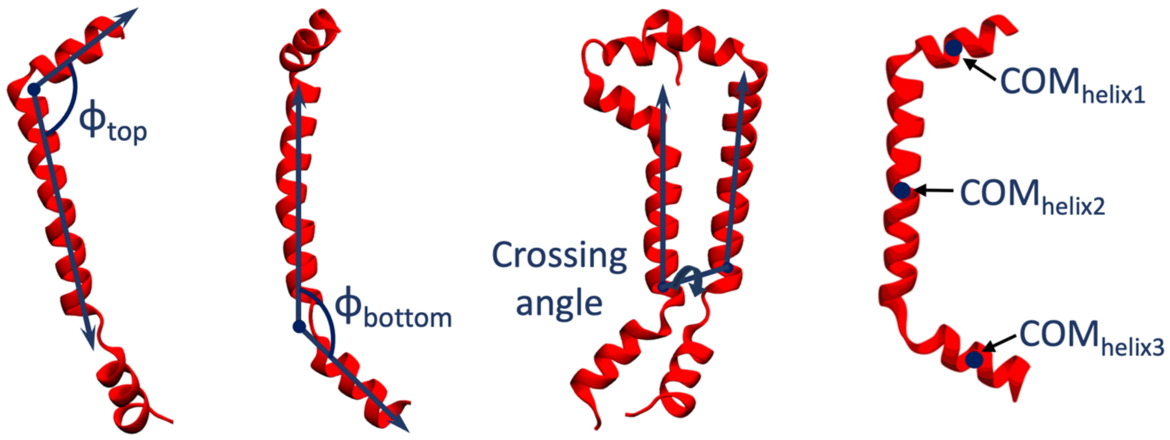


**Supplementary Figure 7:** A total of 36 descriptors were used to characterize a trimer conformation of the MPER-TMD domain. Six descriptors defined the top hinge angles ($\phi_{top}$) and bottom hinge angles ($\phi_{bottom}$) of three chains. Three descriptors defined the crossing angles between the transmembrane helices: helix A and helix B, helix B and helix C, and helix C and helix A. Finally, 27 descriptors captured the inter-chain distances between $COM_{helix1}$, $COM_{helix2}$, and $COM_{helix3}$ for each chain of the MPER-TMD trimer.

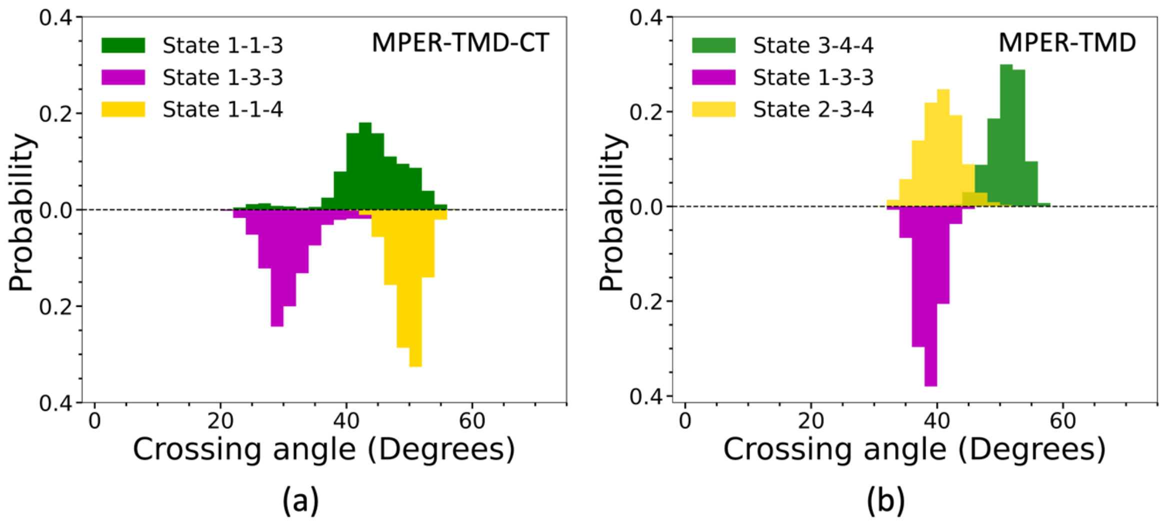


**Supplementary Figure 8:** Distribution of the average crossing angles between transmembrane helices characterizing different conformational states of the (a) MPER-TMD-CT and (b) MPER-TMD trimers.

### Data Availability

Initial and final structures of the asymmetric membrane bilayer containing MPER-TMD-CT/MPER-TMD region of gp41, POPC, LSM, cholesterol, POPE, POPS, and PIP2, protein trajectories, GROMACS input files used to perform equilibration and production runs, and SPIB script are freely available at:

https://github.com/ayan-majumder95/gp41_2025
